# FluoroFate: A generalisable platform for time-resolved single-cell analysis of cell fate enables quantification of cell death dynamics

**DOI:** 10.64898/2026.08.17.745187

**Authors:** Marcus K Preedy, Isobel Taylor-Hearn, Chen Ying, Matthew J Ford, Ian J Jackson, Andrew Gilmore, Vinay Tergoankar, Richard L Mort

## Abstract

Fundamental cellular decisions of life and death are governed by intricate and tightly regulated intracellular signalling pathways that determine whether cells proliferate, enter quiescence, or undergo programmed cell death (apoptosis). Live-cell fluorescence imaging enables these processes to be observed in real time at single-cell resolution, but two problems limit their study. First, existing biosensors do not allow apoptotic status and cell cycle progression to be resolved in tandem within the same cell. Second, interpreting live-cell imaging data is challenging even where multiplex reporters exist, as the biological meaning of fluorescent signals depends on their temporal ordering, and large-scale imaging experiments generate complex, multidimensional data that are difficult to analyse systematically and at scale.

Here we address both problems. We present FluoroFate, a generalisable and user-friendly graphical interface-driven tool for time-resolved single-cell analysis of multiplex live-cell imaging datasets, which integrates existing, robust deep learning-based segmentation, cell tracking, and temporal classification methods to quantify fluorescent reporter dynamics in individual cells across time without the need for specialist computational expertise. Alongside FluoroFate, we develop tricistronic Fluorescent Ubiquitination-based Cell Cycle Indicator (Fucci) and apoptosis biosensors, enabling simultaneous monitoring of cell cycle progression and caspase activation within the same cell.

Applying FluoroFate, we resolve apoptotic and non-apoptotic cell death at the single-cell level based on the temporal ordering of Annexin V and propidium iodide signals, identifying distinct kinetic and phenotypic cell death profiles in response to pharmacological perturbation. We highlight divergent temporal dynamics and modes of cell death between birinapant and cycloheximide treatment, reflecting differences in how TNFα/TNFR1 signalling is disrupted by these agents. At the single-cell level, we uncover parallel, independently regulated death programmes, demonstrating that loss of RIPK1 selectively impairs apoptotic cell death whilst leaving non-apoptotic death largely unaffected.

We then use FluoroFate to analyse timelapse images of our combined Fucci-apoptosis reporters, resolving cell cycle progression and caspase activation within the same cell over time. Together, FluoroFate and our new cell cycle and apoptosis biosensors represent a broadly applicable platform for extracting mechanistic insight from live-cell imaging data.

## INTRODUCTION

Live-cell imaging enables dynamic cellular processes to be quantified at single-cell resolution and is particularly valuable for studying heterogeneous cell-fate decisions. Apoptosis is a fundamental biological process, a regulated form of cell death that is classically described as being immunologically silent (Mustafa et al., 2024). Although heterogeneous apoptotic responses help preserve tissue integrity following damage, they can also produce fractional killing when complete elimination of a cell population is required, such as during cancer treatment (Spencer et al., 2009, Bertaux et al., 2014, Inde et al., 2020). Resolving this heterogeneity requires longitudinal analysis of individual cells, for which fluorescence-based live-cell imaging is particularly well suited.

Fluorescent probes including Annexin V, Apotracker, and propidium iodide (PI) are routinely used to monitor cell death during live-cell imaging (Gelles and Chipuk, 2016, Szalai and Engedal, 2018, Barth et al., 2020). Annexin V and Apotracker bind phosphatidylserine exposed on the outer leaflet of the plasma membrane during apoptosis (Vermes et al., 1995), whereas PI binds nucleic acids following loss of plasma membrane integrity (Riccardi and Nicoletti, 2006). Given that membrane permeabilisation also permits access to inner-leaflet phosphatidylserine, apoptotic and non-apoptotic cell death are most reliably distinguished by the temporal ordering of these signals. Annexin V or Apotracker positivity preceding PI uptake indicates apoptosis, whereas PI uptake without prior phosphatidylserine staining indicates non-apoptotic cell death (**Figure 1**). Time-resolved dual-label imaging can therefore discriminate between these outcomes based on label ordering and time-to-signal (Gelles et al., 2019).

**Figure 1.**
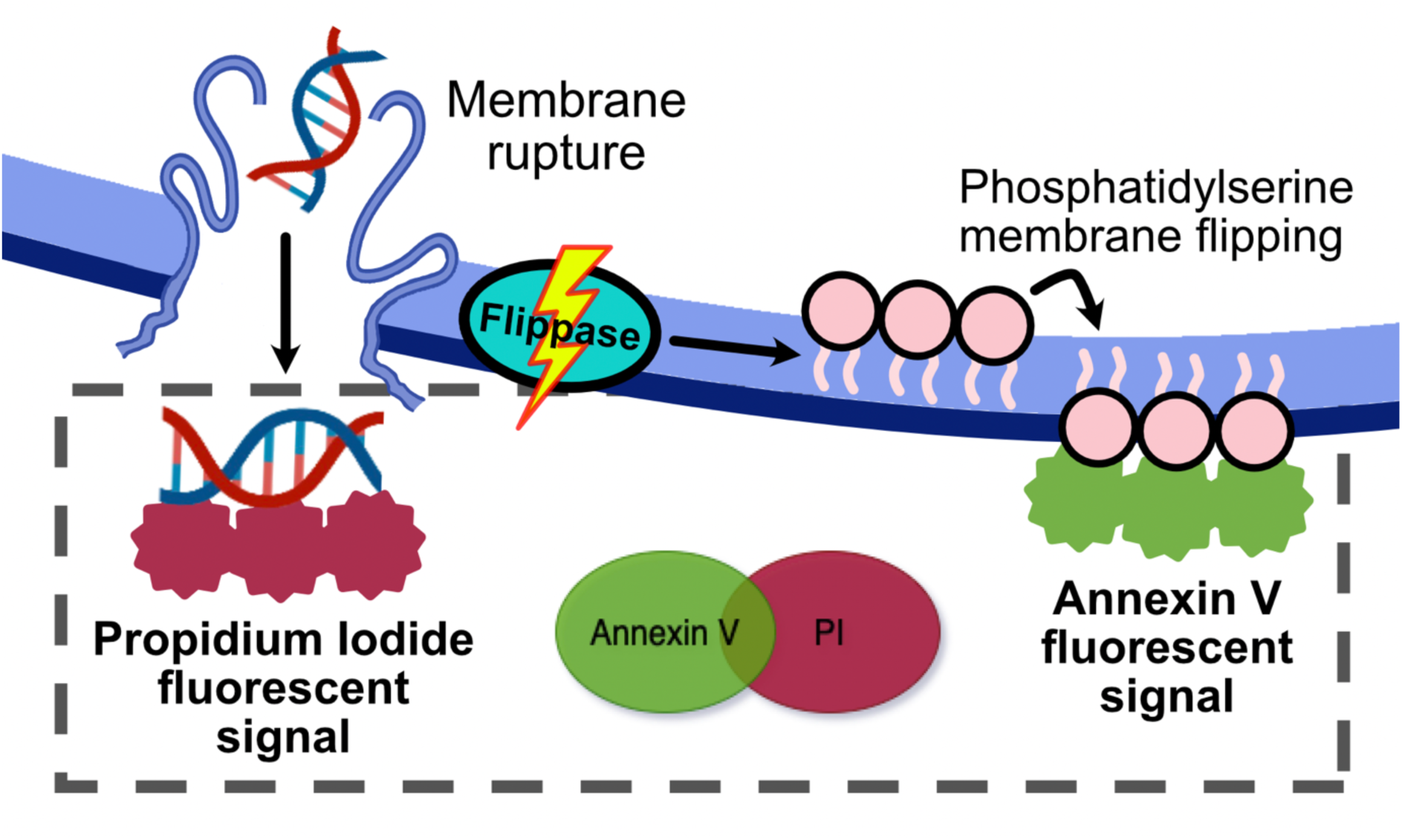
Temporal separation of Annexin V and propidium iodide signals enable discrimination of apoptotic and non-apoptotic cell death. During apoptosis, caspase activation leads to phosphatidylserine externalisation on the outer leaflet of the plasma membrane, enabling binding of Annexin V and generation of a fluorescent signal. At later stages, loss of membrane integrity permits entry of propidium iodide, which intercalates with DNA to produce fluorescence. In contrast, primary loss of membrane integrity in non-apoptotic cell death allows propidium iodide uptake without prior phosphatidylserine exposure. The temporal ordering of Annexin V and propidium iodide signals therefore provide a means to distinguish apoptotic from non-apoptotic cell death at the single-cell level.

A further challenge is determining how apoptotic decision-making relates to the state of an individual cell at the time of treatment. Cell-cycle phase can influence treatment response, yet this relationship is difficult to resolve in unsynchronised populations as cell-cycle progression, stimulus exposure and apoptotic commitment occur dynamically and differ between cells (Beaumont et al., 2016). Genetically encoded reporters provide a means of addressing this problem. The Fluorescent Ubiquitination-based Cell Cycle Indicator (Fucci) system uses cell-cycle-dependent protein degradation to visualise cell-cycle progression in single cells (Sakaue-Sawano et al., 2008), with subsequent systems providing more precise resolution of cell-cycle stages (Sakaue-Sawano et al., 2017). Fucci models have also been developed for whole-organism imaging in species including mouse and chicken (Sakaue-Sawano et al., 2008, Mort et al., 2014, Ford et al., 2018, Sudderick et al., 2026). However, simultaneous analysis of cell-cycle state and apoptotic commitment requires reporters that combine cell-cycle information with a direct readout of caspase activation.

Large live-cell imaging datasets also require robust computational analysis. Recent advances in cellular segmentation (Stringer et al., 2021, Pachitariu et al., 2025, Archit et al., 2025) and automated cell tracking (Ershov et al., 2022, Bragantini et al., 2025) have enabled high-throughput extraction of single-cell fluorescence trajectories. However, converting these trajectories into biologically meaningful classifications still requires accessible tools that integrate segmentation, tracking, and temporal interpretation of multiple fluorescent reporters.

Here, we present FluoroFate (https://github.com/isobelth/FluoroFate), a user-friendly graphical interface that combines Cellpose (Pachitariu et al., 2025), TrackMate (Ershov et al., 2022), and temporal classification of fluorescent signals. FluoroFate integrates open-source tools implemented across different software environments, including Python and Fiji (Schindelin et al., 2012), into a single platform, providing a streamlined workflow for image analysis. We also develop two Fucci-apoptosis biosensor systems that enable simultaneous monitoring of cell-cycle progression and caspase activation in individual cells. Together, these biosensors and FluoroFate allow cell-cycle phase to be assigned at the time of apoptotic commitment and enable apoptotic and non-apoptotic cell death to be distinguished from the temporal ordering of fluorescent markers.

## RESULTS

### FluoroFate: a user-friendly analysis tool for single cells in fluorescent timelapse imaging datasets

Advances in cellular segmentation and tracking have facilitated automated single-cell analysis of live-cell imaging data. Here, we integrate Cellpose (Pachitariu et al., 2025) and TrackMate (Ershov et al., 2022) within the FluoroFate graphical interface to segment and track cells from brightfield images. Downstream of segmentation and tracking, custom algorithms are applied to classify cells based on the timing and intensity of fluorescent signals. Users can analyse up to three fluorescent channels and define intensity thresholds for each (see Materials and Methods). FluoroFate supports two complementary analysis modes: **persistent** and **dynamic** analysis. Persistent analysis is primarily designed for cell death assays, in which cells are irreversibly classified once a fluorescence threshold is exceeded. This enables classification of distinct cell death modalities based on the temporal ordering of fluorescent signal acquisition. In contrast, dynamic analysis re-evaluates fluorescence intensity at each time point, allowing cell identities to change dynamically across frames. This mode is particularly suited to reporters with dynamic expression patterns, such as Fucci-based cell cycle indicators. **Figure 2** provides an overview of this workflow.

**Figure 2.**
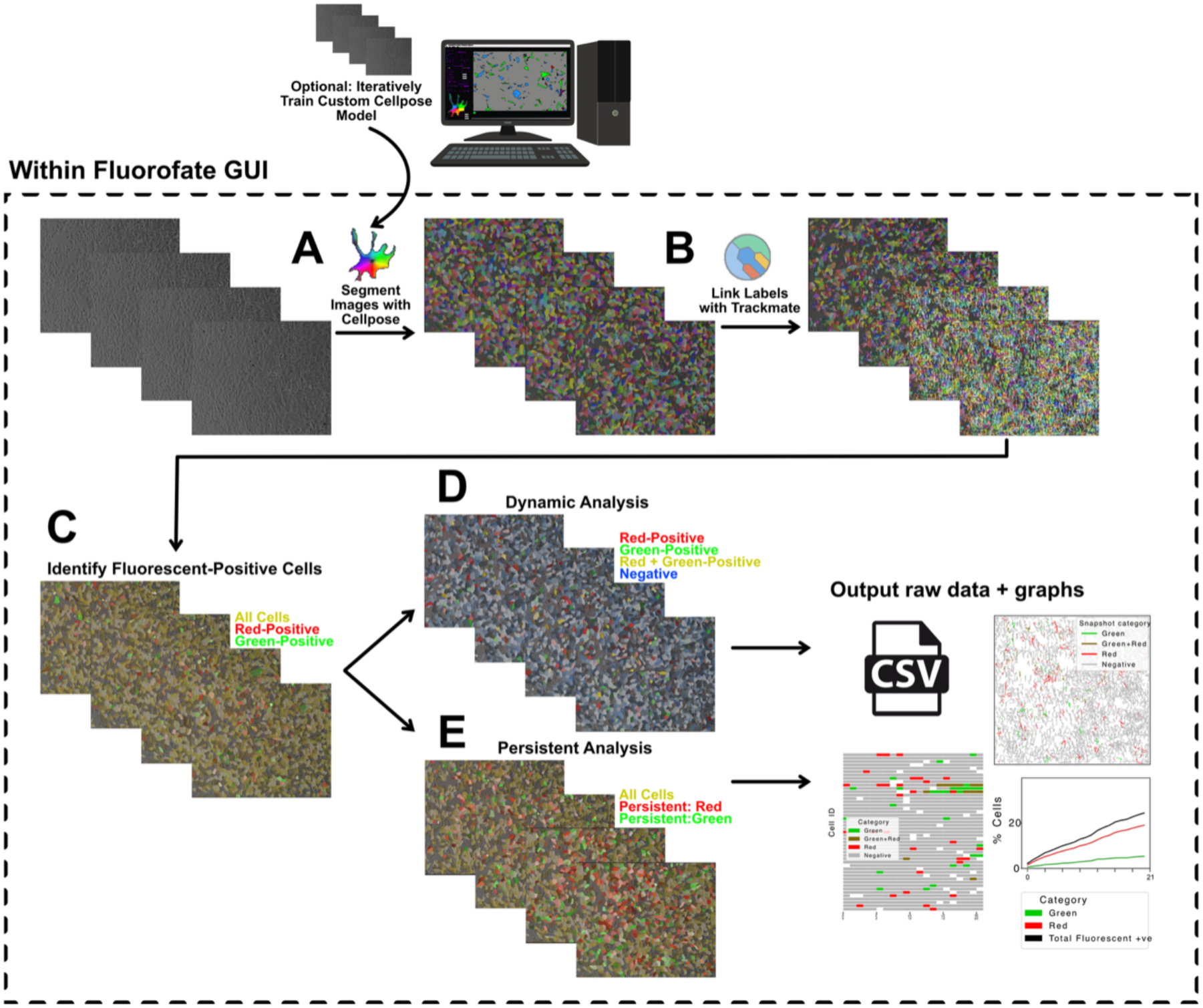
FluoroFate workflow for integrated single-cell timelapse analysis. Multi-channel timelapse microscopy data are analysed within the FluoroFate graphical user interface, which brings segmentation, tracking, fluorescence quantification, and fate classification into a single workflow. (**A**) Cells are segmented from the brightfield channel using Cellpose. (**B**) Segmented objects are then linked across frames using TrackMate to generate consistent single-cell tracks over time. (**C**) Fluorescence positivity is assigned to each tracked cell for up to three user-defined channels following per-channel thresholding. The resulting data can be analysed in two complementary modes: (**D**) **dynamic analysis**, which classifies cells according to their fluorescence state at each individual timepoint, and [**E**] **persistent analysis**, which assigns cells according to the first fluorophore they persistently express, thereby capturing the temporal order of reporter activation. By integrating Cellpose- and TrackMate-based analysis within a single GUI, FluoroFate removes the need to move between ImageJ, Python scripts, and separate visualisation tools, enabling straightforward and reproducible end-to-end analysis. Both modes export cell-level CSV outputs together with graphical summaries, including spatial track plots, cell-timeline visualisations, and percentage-over-time trajectories.

To facilitate accessible and streamlined analysis of live-cell imaging datasets, we developed FluoroFate as a graphical user interface (GUI) that integrates segmentation, tracking, and temporal classification within a single platform. Built using Napari (Chi-Li and Clack, 2022) and magicgui (https://pypi.org/project/magicgui/), the interface allows users to perform the full analysis workflow without the need for coding or switching between multiple software environments. By running Fiji/ImageJ in headless mode, users can access TrackMate from within the Python environment. Key parameters, including image input (tiff format), channel assignment, fluorescence thresholds, and TrackMate settings can be adjusted directly within the GUI. Cellular segmentation and tracking are also performed in the GUI. The Napari viewer provides an interactive workspace for visualising raw images alongside segmentation masks, tracked cells, and classified outputs in real time (**Figure 3**). Together, these features enable intuitive, high-throughput analysis of complex live-cell imaging datasets without the need for programming expertise. The GUI has been designed and tested for MacOS, Linux, and Windows operating systems.

**Figure 3.**
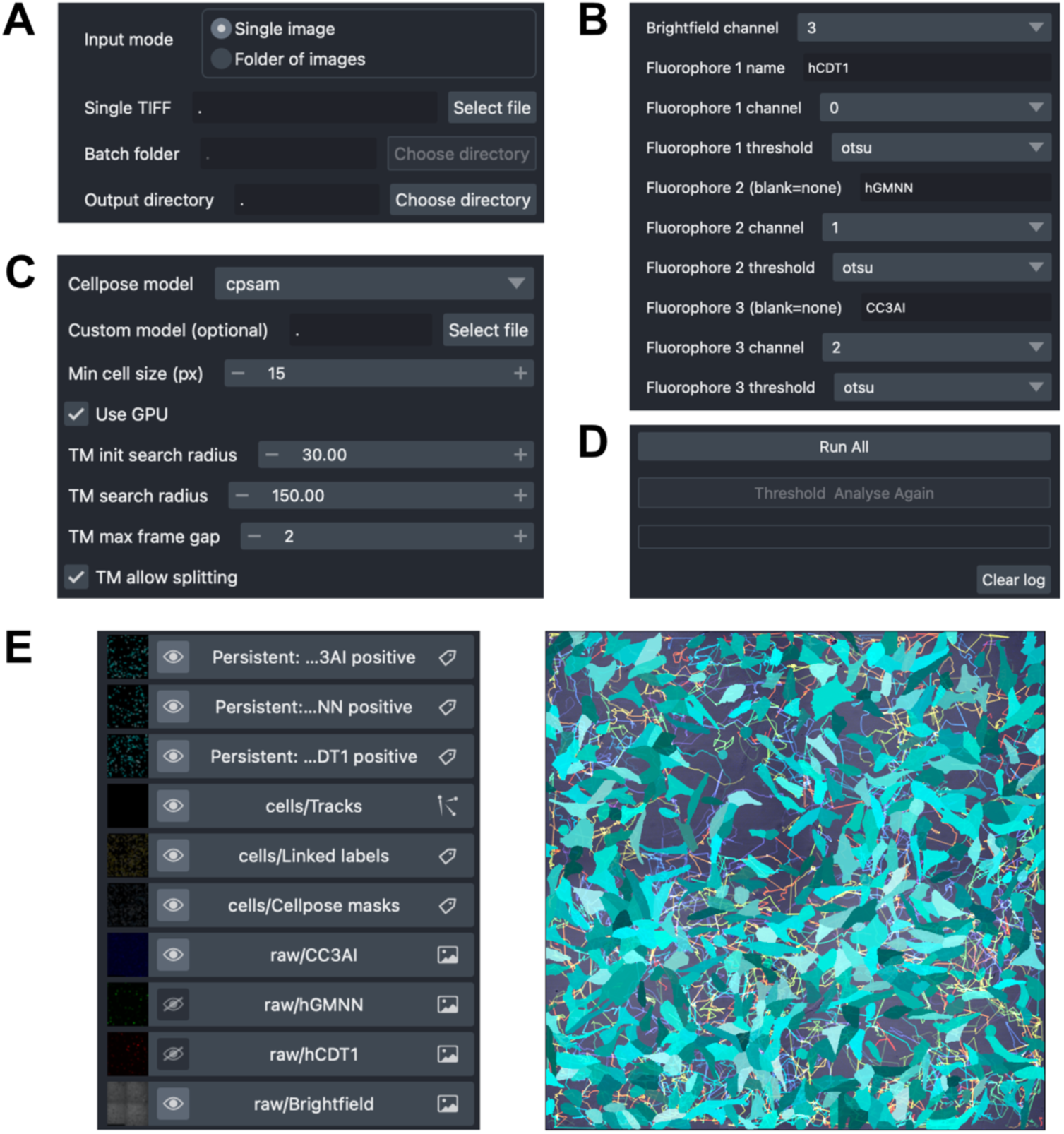
FluoroFate graphical user interface built with napari and magicgui. Screenshot of the FluoroFate GUI, built with Napari and magicgui, that enables integrated time-lapse microscopy analysis. Docked magicgui panels provide access to (**A**) file selection, (**B**) channel assignment and thresholding settings (**C**) Cellpose model choice, and TrackMate tracking parameters. (**D**) A central “Run All” panel enables users to execute segmentation, tracking, persistent analysis and dynamic analysis directly from the same interface. [**E**] The Napari viewer displays the brightfield image, fluorescence channels, and output Cellpose segmentation masks, TrackMate-linked labels, and fluorescence-positive cells in a single interactive workspace. By combining established segmentation and tracking tools within one accessible GUI, FluoroFate lowers the technical barrier to rigorous single-cell time-lapse analysis and avoids the need to switch between ImageJ/Fiji, Python, and separate visualisation environments during analysis.

### FluoroFate analysis reveals distinct TNFα-induced cell death dynamics under birinapant and cycloheximide treatment

In response to TNFα stimulation, TNFR1 signalling is typically initiated through the formation of complex I, which promotes activation of NF-κB and MAPK pathways and drives the expression of pro-survival and inflammatory genes (Tian et al., 2005, Varfolomeev et al., 2007). Disruption of this signalling can shift the cellular response towards cell death through formation of cytosolic complex II, resulting in caspase activation and apoptosis (Micheau and Tschopp, 2003). Pharmacological agents can perturb this pathway at distinct stages. The SMAC mimetic birinapant promotes degradation of cIAP1/2, destabilising complex I and favouring complex II formation, thereby sensitising cells to TNFα-induced apoptosis (Li et al., 2004, Vince et al., 2007, Wang et al., 2008, Feltham et al., 2011, Benetatos et al., 2014). In contrast, cycloheximide inhibits protein synthesis, preventing the accumulation of anti-apoptotic factors downstream of TNFR1 signalling (Wang et al., 2008) (**Figure 4A**).

**Figure 4.**
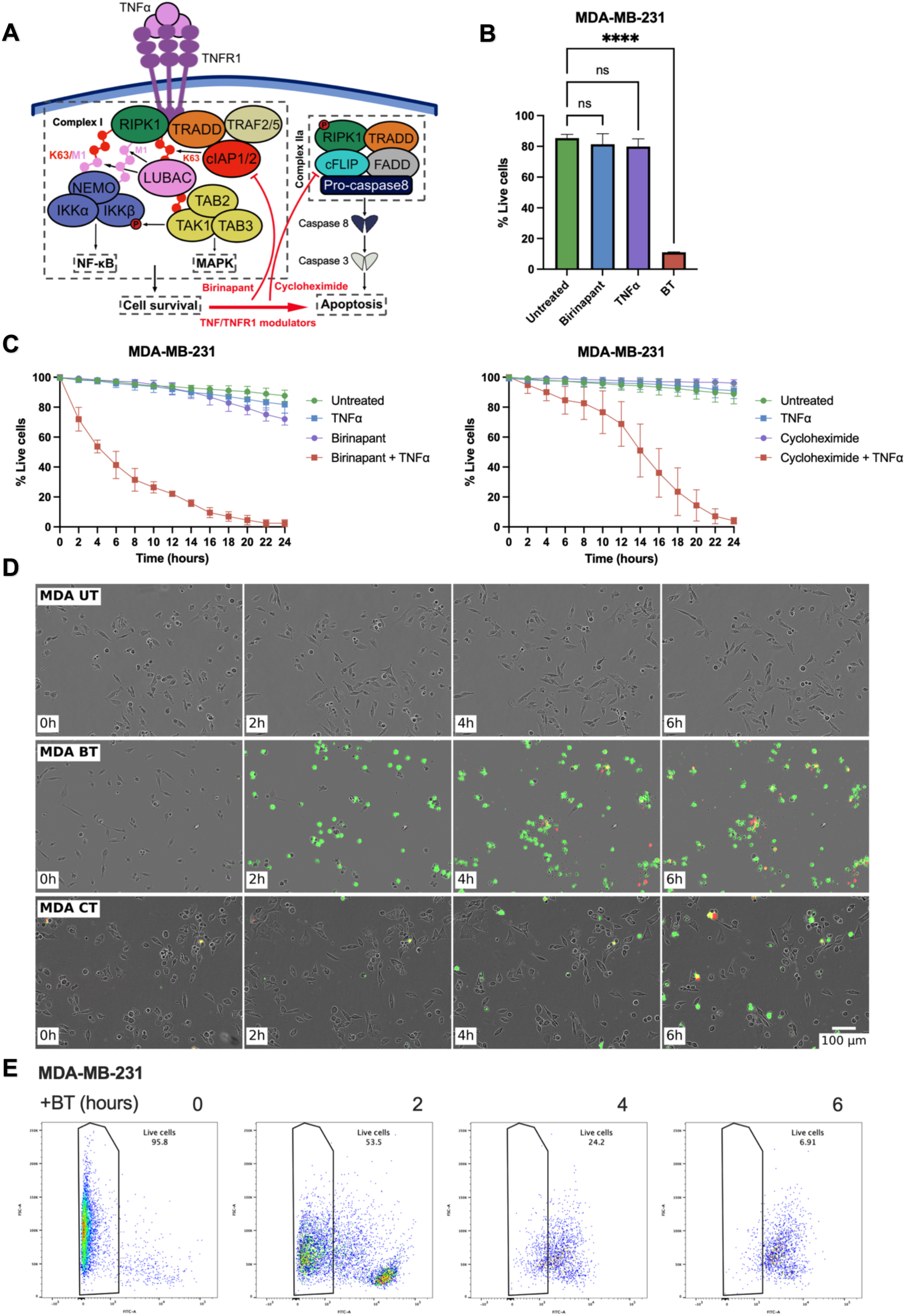
Distinct TNFα-induced cell death dynamics under birinapant and cycloheximide treatment. (**A**) Schematic overview of TNFα/TNFR1 signalling. Engagement of TNFR1 promotes formation of membrane-associated complex I, leading to activation of NF-κB and MAPK signalling pathways and induction of pro-survival gene expression. Disruption of this pathway can result in formation of cytosolic complex IIa, leading to caspase activation and apoptosis. Birinapant promotes degradation of cIAP1/2, destabilising complex I and favouring early complex II formation, whereas cycloheximide inhibits protein synthesis and prevents accumulation of pro-survival factors downstream of TNFR1 signalling. (**B**) Quantification of live cells following treatment with TNFα, birinapant, or their combination (BT), assessed by flow cytometry in MDA-MB-231 cells. Cells were stained with Annexin V and propidium iodide (PI), and live cells were defined as Annexin V-negative/PI-negative. Data represent mean ± SD from three independent biological replicates. (**C**) Time-resolved analysis of cell viability following treatment with TNFα in combination with birinapant (**left**) or cycloheximide (**right**). Live-cell percentages were quantified from IncuCyte live-cell imaging using FluoroFate. BT treatment induces rapid and near-complete loss of viability, whereas CT treatment results in delayed but progressive cell death, demonstrating distinct temporal dynamics depending on the mode of TNFα pathway perturbation. (**D**) Representative live-cell imaging of MDA-MB-231 cells over the first 6 hours following treatment with untreated control (UT), BT, or CT. Annexin V-positive cells (green) and PI-positive cells (red) are shown overlaid on brightfield images. BT treatment results in rapid and widespread Annexin V staining, whereas CT treatment shows delayed onset of cell death. Scale bar, 100 μm. [**E**] Flow cytometry analysis of Annexin V staining over the first 6 hours following BT treatment. Representative dot plots show progressive increase in Annexin V-positive cells over time, confirming rapid induction of apoptosis at the single-cell level. Statistical significance was determined using one-way ANOVA with Dunnett’s post-hoc test, with **** indicating P < 0.0001 and ns indicating non-significant differences.

Given these distinct mechanisms, we sought to determine whether birinapant and cycloheximide produce different cell death dynamics at both the population and single-cell level. To address this, we compared the temporal progression and phenotypic characteristics of TNFα-induced cell death following treatment with birinapant plus TNFα (BT) or cycloheximide plus TNFα (CT). Timelapse live-cell imaging of treated cells was analysed using FluoroFate in persistent mode. In MDA-MB-231 (triple-negative breast cancer) cells, BT treatment resulted in rapid and near-complete cell death (**Figure 4C left**), whereas CT induced a delayed but progressive loss of viability (**Figure 4C right**). Induction of cell death in response to BT treatment was also assessed via flow cytometry (**Figures 4B, E**).

### FluoroFate analysis reveals RIPK1-dependent apoptotic and non-apoptotic cell death dynamics

RIPK1 is a key regulator of TNFα-induced cell fate, with its post-translational modification and caspase 8-mediated cleavage influencing the balance between survival, apoptosis, and necroptosis (Feltham and Silke, 2017, Newton et al., 2019, Wang et al., 2021, Ju et al., 2022). We therefore used FluoroFate to determine how RIPK1 loss affects the kinetics and mode of TNFα-induced cell death at the single-cell level. To address this, we compared responses to BT treatment in SKOV3 (human ovarian cancer), WT, and RIPK1 KO cells using FluoroFate in persistent mode (**Figure 5**). In parallel, we assessed the relative contributions of apoptotic and non-apoptotic cell death in MDA-MB-231 cells following BT or CT treatment, revealing a more robust and rapid induction of apoptosis in response to BT than CT (**Figure 5C**). Induction of apoptosis under BT treatment was confirmed by western blot analysis of caspase activation and PARP cleavage (**Figure 5B**). RIPK1 KO cells exhibited marked resistance to BT-induced cell death compared to WT cells (**Figures 5D, E**). Interestingly, loss of RIPK1 selectively abrogated apoptotic cell death, whilst non-apoptotic cell death remained largely unchanged (**Figure 5F**), indicating that these modes of cell death can be independently regulated at the single-cell level. These results highlight the ability of our platform to resolve distinct, independently regulated modes of cell death within heterogeneous cell populations.

**Figure 5.**
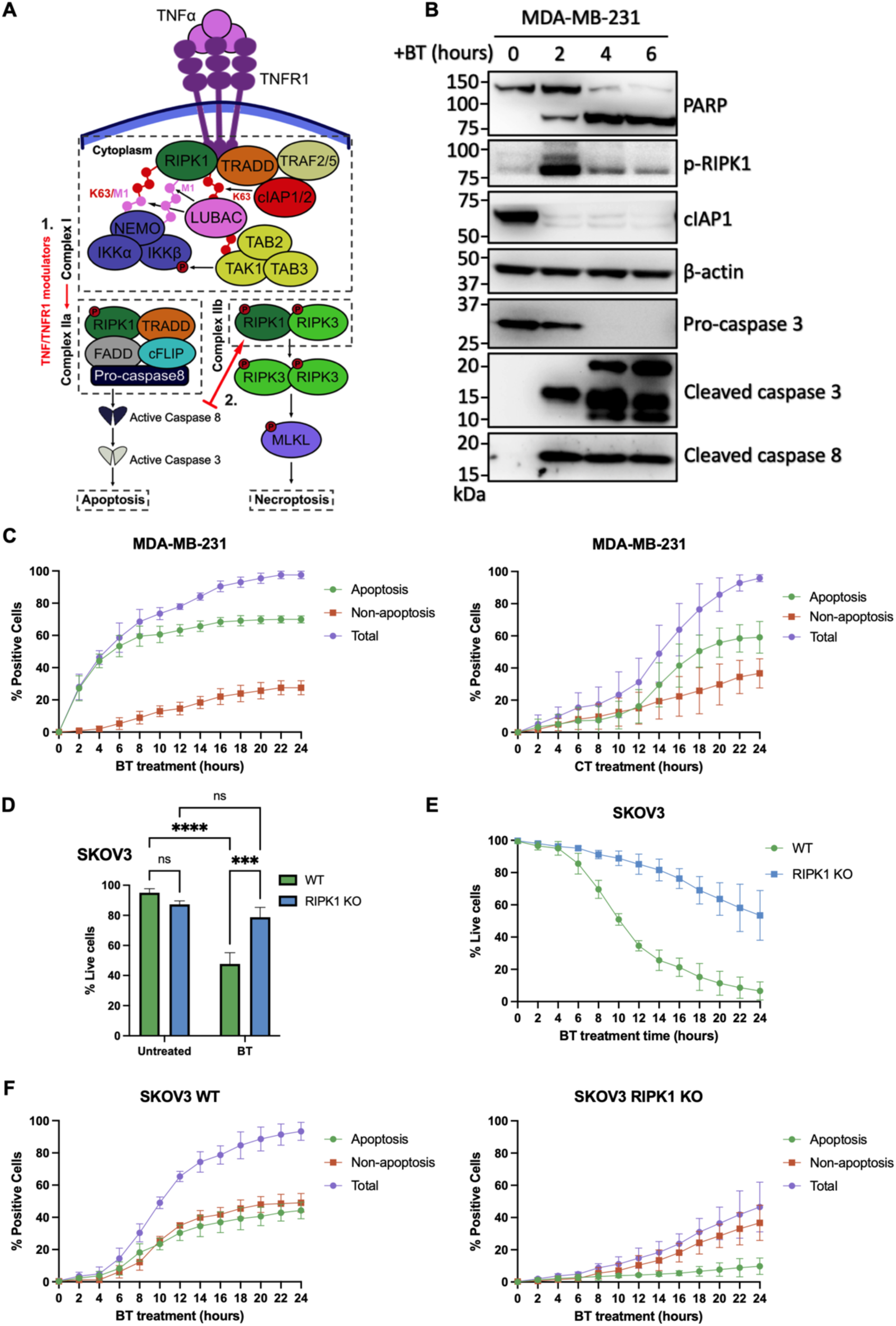
Single-cell analysis reveals RIPK1-dependent apoptotic and non-apoptotic cell death dynamics following TNFα stimulation. (**A**) Schematic overview of TNFα/TNFR1 signalling, illustrating that disruption of complex IIa can promote formation of cytosolic complex IIb, from which signalling through RIPK1/RIPK3/MLKL can drive non-apoptotic cell death. (**B**) Western blot analysis of apoptotic signalling following BT treatment in MDA-MB-231 cells over the indicated time course. Cleavage of PARP, caspase 3, and caspase 8 confirms induction of apoptosis. Autophosphorylation of RIPK1 (p-RIPK1) at Ser166 and reduction of cIAP1 levels demonstrate modulation of TNFα/TNFR1 signalling. Β-actin serves as a loading control. (**C**) FluoroFate analysis of cell death dynamics in MDA-MB-231 cells following treatment with BT (**left**) or CT (**right**). The proportion of cells undergoing apoptosis (Annexin V-positive prior to PI), non-apoptotic cell death (PI-positive prior to or concurrent with Annexin V), and total cell death is shown. BT induces rapid and predominantly apoptotic cell death, whereas CT results in delayed and more heterogeneous cell death dynamics. (**D**) Quantification of live cells in SKOV3 WT and RIPK1 KO cells following BT treatment, assessed by flow cytometry. Cells were stained with Annexin V-FITC and PI, and live cells were defined as Annexin V-negative/PI-negative. RIPK1 KO cells exhibit increased resistance to BT-induced cell death compared to WT cells. [**E**] FluoroFate analysis of cell viability in SKOV3 WT and RIPK1 KO cells following BT treatment. Loss of RIPK1 delays and reduces overall cell death kinetics. (**F**) Single-cell classification of apoptotic and non-apoptotic cell death in SKOV3 WT (**left**) and RIPK1 KO (**right**) cells following BT treatment. Loss of RIPK1 selectively reduces apoptotic cell death, whilst non-apoptotic cell death remains largely unchanged, indicating that distinct modes of cell death are independently regulated at the single-cell level. Data represent mean ± SD from three biological replicates. Statistical significance was determined using two-way ANOVA followed by Fisher’s LSD post-hoc test (***P ≤ 0.0002, ****P < 0.0001, ns, not significant).

### Development of Fucci-apoptosis reporter systems enables simultaneous monitoring of cell cycle progression and caspase activation with FluoroFate

Cell-fate responses to external stimuli can be influenced by the cell-cycle stage at which the signal is received, contributing to phase-specific differences in drug sensitivity and resistance (Beaumont et al., 2016). To extend our analysis platform to the simultaneous measurement of proliferation and cell death, we developed and evaluated dual-function reporter constructs combining fluorescent cell cycle indicators (tFucci2(SA)) (Mort et al., 2014) with caspase-activated fluorescent apoptosis reporters. The Fucci system exploits cell cycle-dependent ubiquitin-mediated proteasomal degradation of fluorescent reporters fused to CDT1 and geminin degrons (Sakaue-Sawano et al., 2008). The CDT1 reporter accumulates during G1 and is degraded at G1/S, whereas the geminin reporter accumulates during S/G2 and is degraded at mitotic exit (**Figure 6A**). To monitor apoptosis alongside cell cycle stage, two candidate caspase-3-responsive reporters were assessed: a cleavage-activated fluorophore based on a quenched GFP design (CA-Cerulean), and a cyclisation-based reporter (CC3AI). The CA-Cerulean reporter is derived from a previously described caspase-activatable GFP construct, in which a quenching peptide prevents fluorophore maturation until cleavage at a DEVD caspase recognition sequence removes the quenching peptide, allowing correct folding of the fluorescent protein (Nicholls et al., 2011). We substituted EGFP for Cerulean to achieve optimal spectral separation from tFucci2(SA). In parallel, the CC3AI reporter is a circularly permuted Cerulean variant whose new termini are joined by a split Npu DnaE intein, cyclising the protein into a constitutively non-fluorescent resting state; caspase-mediated cleavage of an internal DEVDG linker opens this loop, converting the cyclised protein into a linear form that folds correctly and restores fluorescence (Zhang et al., 2013). Both reporters were engineered into tricistronic constructs by fusing to tFucci2(SA) using a P2A self-cleaving peptide, enabling simultaneous expression of cell cycle and apoptosis reporters within the same cell.

**Figure 6.**
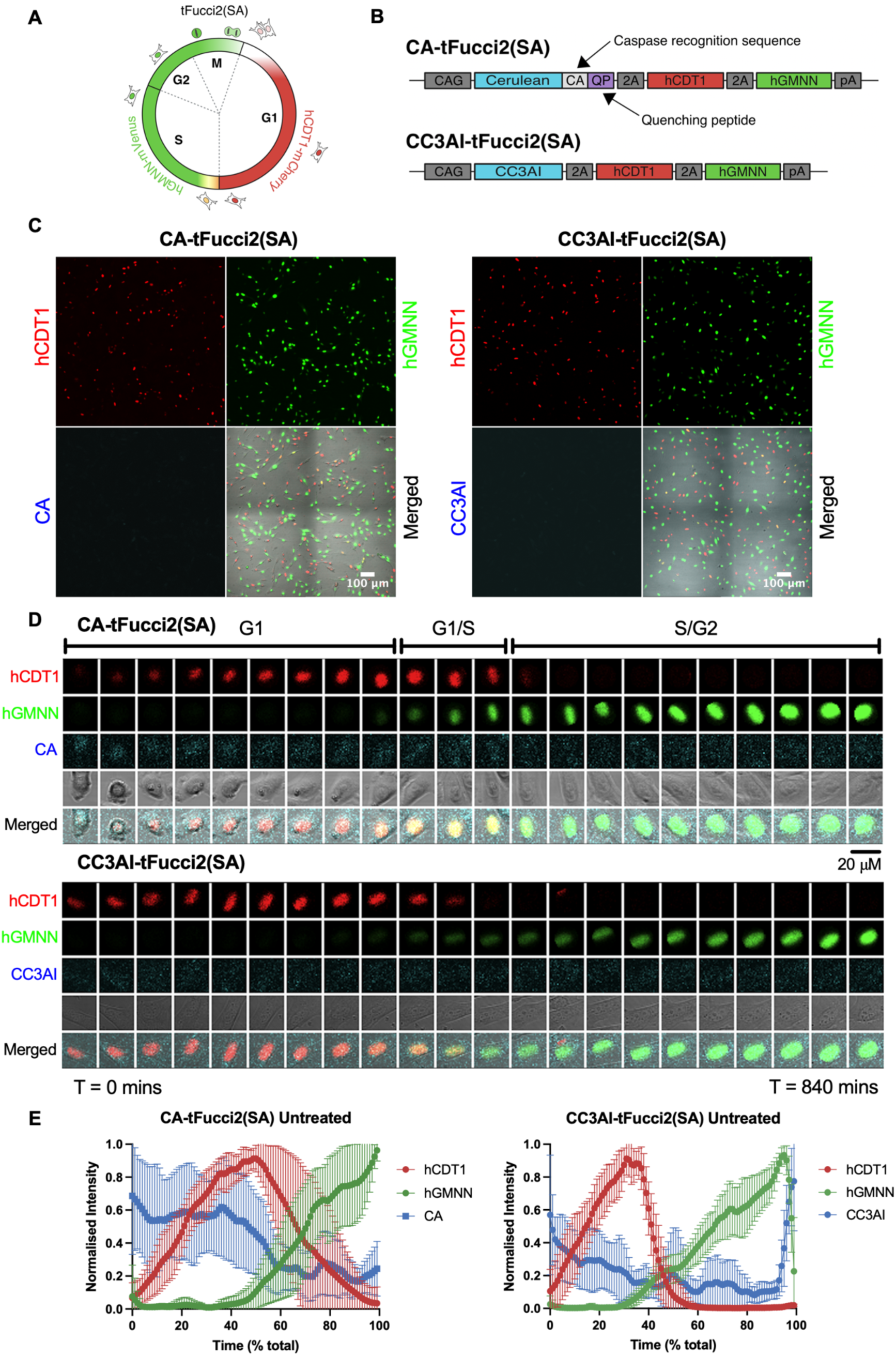
CA-tFucci2(SA) and CC3AI-tFucci2(SA) reporters show normal cell cycle progression and regulation of Fucci biosensors. (**A**) Diagram of the tFucci2(SA) reporter system used to classify cell cycle stage through phase-specific fluorescent expression. Cells in G1 are labelled red through expression of mCherry-hCDT1, and cells in S/G2/M are labelled green through expression of mVenus-hGeminin. Transitional states are represented by dual-positive fluorescence. (**B**) Overview of the tricistronic CA-tFucci2(SA) and CC3AI-tFucci2(SA) reporter constructs. Both constructs combine a caspase-activatable Cerulean apoptosis reporter with the tFucci2(SA) cell cycle reporters using self-cleaving P2A peptides. In the CA-tFucci2(SA) construct, Cerulean fluorescence is suppressed by a quenching peptide linked via a caspase recognition sequence, whereas CC3AI-tFucci2(SA) contains a split-fluorophore apoptotic reporter activated following caspase cleavage. (**C**) Representative microscopy images of CA-tFucci2(SA)- (**left**) and CC3AI-tFucci2(SA)- (**right**) expressing cells in untreated conditions. (**D**) Timelapse montage of a single cell expressing CA-tFucci2(SA) (**top**) or CC3AI-tFucci2(SA) (**bottom**), highlighting normal cell cycle progression and Fucci biosensor expression. [**E**] Mean Fucci biosensor dynamics across a complete cell cycle (n = 10 tracked cells). Individual cell traces were interpolated and aggregated using FucciTools (Van Kerckvoorde et al., 2021).

### CA-tFucci2(SA) and CC3AI-tFucci2(SA) reporters show normal cell cycle progression and regulation of Fucci biosensors

Stable NIH 3T3 cell lines expressing either CA-tFucci2(SA) or CC3AI-tFucci2(SA) (**Figure 6B**) were generated using the Flp/In system and imaged using timelapse microscopy. Under basal conditions, Fucci reporters displayed correct localisation and cell-cycle phase specificity, with CDT1 reporter signal increasing during G1 and declining as geminin reporter signal increased upon entry into S phase. Geminin signal continued to increase throughout G2 (**Figures 6D, E**). CA-Cerulean and CC3AI signals remained low throughout the cell cycle (**Figure 6D**).

### FluoroFate simultaneously monitors cell-cycle dynamics and caspase activation in CA-tFucci2(SA)-and CC3AI-tFucci2(SA)-expressing cells

Timelapse images of untreated cells expressing CA-tFucci2(SA) or CC3AI-tFucci2(SA) were analysed using FluoroFate. Dynamic analysis was performed to quantify the percentage of CDT1- and geminin-positive cells per frame. Interestingly, cycling of the Fucci reporters was detectable at the population level (**Figure 7A**). To assess Cerulean fluorescence during apoptosis, cells were treated with staurosporine. Following treatment, cycling of the Fucci reporters ceased, with cell cycle states remaining effectively static during the rapid and widespread induction of apoptosis (**Figures 7B, D**). Both constructs showed increased Cerulean fluorescence, indicating caspase activation; however, CA-Cerulean exhibited a greater dynamic range (∼1.9-fold) than CC3AI (∼1.5-fold) (**Figure 7C**).

**Figure 7.**
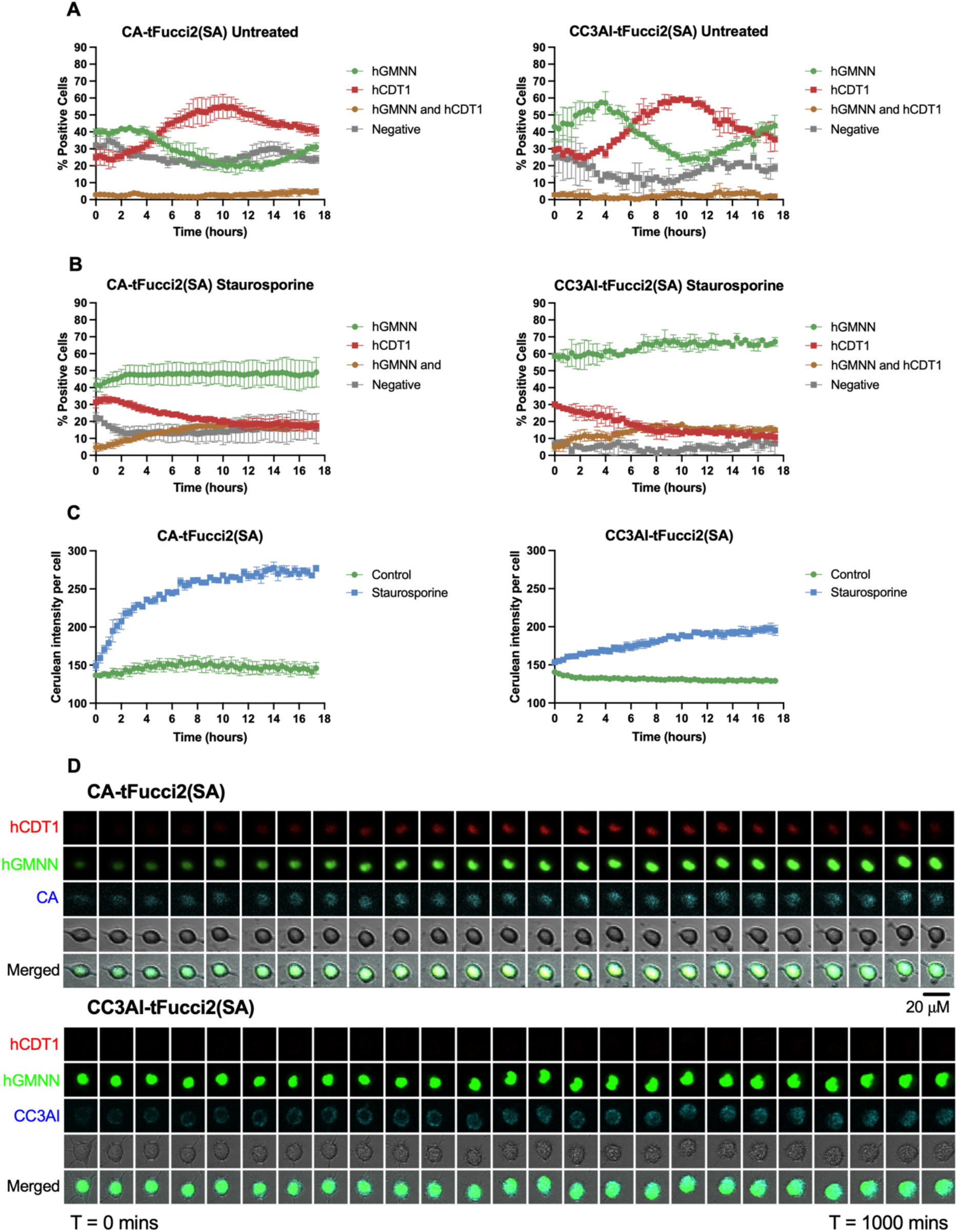
FluoroFate enables analysis of combined Fucci-apoptosis reporter systems for simultaneous monitoring of cell cycle progression and caspase activation. Dynamic analysis of Fucci fluorescence states over time in (**A**) untreated and (**B**) staurosporine-treated cells expressing CA-tFucci2(SA) (**left**) or CC3AI-tFucci2(SA) (**right**). Cells were classified as hGMNN-positive, hCDT1-positive, dual-positive, or negative at each time point using FluoroFate dynamic analysis. (**C**) Quantification of mean Cerulean fluorescence intensity per unit cell area over time in CA-tFucci2(SA)-(**left**) and CC3AI-tFucci2(SA)- (**right**) expressing cells following staurosporine treatment. Fluorescence intensity was measured for individual cells, normalised to cell area, and averaged across all cells at each frame. Both reporters exhibited increased Cerulean fluorescence in response to apoptosis induction, with CA-tFucci2(SA) producing a larger dynamic fluorescence increase relative to untreated controls. (**D**) Timelapse montage of a single cell expressing CA-tFucci2(SA) (**top**) or CC3AI-tFucci2(SA) (**bottom**) in response to staurosporine treatment. Data represent mean ± SD from three biological (CA- tFucci2(SA)) or two biological (CC3AI-tFucci2(SA)) replicates.

## DISCUSSION

FluoroFate enables time-resolved, single-cell analysis of fluorescent live-cell imaging datasets and, in the context of cell death assays, can distinguish apoptotic from non-apoptotic death based on signal ordering rather than endpoint staining alone. This is an important advance, as many cell death markers are not mechanism-specific when interpreted in isolation. Annexin V and related phosphatidylserine-binding probes are frequently used as apoptotic markers, yet loss of membrane integrity exposes inner leaflet phosphatidylserine and generates fluorescent signal in cells undergoing non-apoptotic forms of death (Vermes et al., 1995, Waring et al., 1999). By incorporating temporal information, FluoroFate allows these signals to be decoded in the sequence in which they arise, providing a more informative view of cell death mechanisms at the single-cell level. This temporal classification is particularly valuable in the context of pharmacological perturbation, where the kinetics and mechanism of death reflect how intracellular signalling pathways have been disrupted (Kreuz et al., 2001, Vince et al., 2007, Benetatos et al., 2014). Smac mimetics have attracted considerable interest as anti-cancer agents, and multiple compounds, including birinapant, APG-1387, and HGS1029, have been evaluated clinically (Morrish et al., 2020). As these agents are being trialled in the clinic, it is essential to understand how they modulate intracellular signalling pathways at a mechanistic level.

Using this approach, we show that birinapant plus TNFα and cycloheximide plus TNFα do not simply induce the same endpoint at different rates but instead generate clearly distinct cell death profiles. BT induced more rapid and robust apoptosis, whereas CT produced delayed and more heterogeneous death dynamics, with a greater relative contribution from non-apoptotic cell death. These differences are consistent with the idea that disruption of TNFα/TNFR1 signalling at different stages produces distinct temporal and phenotypic outcomes and likely reflects the more targeted mechanism of action of birinapant compared to the global inhibition of protein synthesis induced by cycloheximide (Kreuz et al., 2001, Vince et al., 2007, Benetatos et al., 2014). Importantly, the ability to resolve these dynamics at the single-cell level avoids masking effects inherent to population-averaged measurements (Spencer and Sorger, 2011). A further strength of the platform is that it can reveal parallel death processes within the same population. In SKOV3 cells, loss of RIPK1 markedly reduced the apoptotic component of BT-induced death, whilst the non-apoptotic component appeared largely independent of RIPK1. This suggests that at least two separable death programmes contribute to the overall response: one that is strongly RIPK1-dependent and predominantly apoptotic, and another that is RIPK1-independent. One possibility is that the RIPK1-independent component reflects necroptotic or necroptosis-like signalling, as non-canonical routes to necroptosis have been described in some contexts (Wang et al., 2019, Wu et al., 2014). However, this remains speculative and would require additional validation, for example by assessing phosphorylation of RIPK3 and MLKL, and by testing sensitivity to pharmacological inhibitors of these proteins (Kaiser et al., 2013, Dai et al., 2025). At present, our data support the conclusion that FluoroFate can resolve distinct, independently regulated cell death modalities, but do not yet definitively assign the RIPK1-independent component to a specific molecular pathway.

Interpretation of long-term birinapant responses also requires caution. Birinapant is well established as a potent antagonist of cIAP1/2, and cIAP depletion can stabilise NF-κB-inducing kinase (NIK), promoting activation of non-canonical NF-κB signalling (Vince et al., 2007, Varfolomeev et al., 2007). Whilst complex I-mediated activation of canonical NF-κB signalling occurs within minutes of TNFR1 ligation, activation of the non-canonical NF-κB pathway takes several hours and requires new protein synthesis (Zarnegar et al., 2008). Thus, although the early response to BT is likely dominated by acute perturbation of TNFα/TNFR1 signalling, prolonged treatment may influence broader signalling networks beyond the initial complex I/complex II decision. These considerations highlight the value of coupling single-cell temporal analysis with orthogonal biochemical validation, as performed here using western blotting and flow cytometry.

Beyond cell death assays, FluoroFate is designed as a generalisable platform for the anlysis of multiplex live-cell imaging. We show that it can classify cell cycle stage in Fucci datasets and can be paired with combined Fucci-apoptosis reporter systems to simultaneously monitor cell cycle progression and caspase activation within the same cell. This expands the utility of the platform beyond cell death classification and demonstrates that it can analyse datasets in which reporter expression changes dynamically across time. In this respect, FluoroFate is well suited not only to questions of cell death mechanisms, but more broadly to problems in which cellular state transitions must be resolved at single-cell resolution (Sakaue-Sawano et al., 2017). The platform also has practical advantages: by integrating segmentation, tracking, and temporal classification into a user-friendly workflow, FluoroFate substantially reduces the time required to analyse large imaging datasets and makes high-throughput live-cell analysis more accessible. This is particularly relevant for pharmacological screening and for studies of heterogeneous responses, where manual analysis rapidly becomes impractical (Isherwood et al., 2011). The novelty of the platform therefore lies not in the invention of entirely new segmentation or tracking algorithms, but in the combination of established high-performing tools into a coherent, accessible workflow designed specifically for time-resolved interpretation of fluorescent live-cell imaging data.

A potential future direction is extension towards label-free analysis. Recent work has highlighted the growing potential of label-free live-cell imaging approaches, including brightfield-based analysis and quantitative phase imaging, for long-term cell monitoring and cell death detection (Chen et al., 2023, Chen et al., 2024). Such approaches may be particularly valuable in settings where exogenous fluorescent reporters are undesirable, limited by spectral complexity, or difficult to implement *in vivo*. In that context, FluoroFate provides a useful conceptual framework: although developed here for fluorescent reporter-based datasets, its core logic of time-resolved single-cell classification could in principle be extended to label-free readouts as these methods mature.

## CONCLUSIONS AND OUTLOOK

Here we present FluoroFate, a code-free GUI that automates single-cell image analysis using dynamic and persistent fluorescence measurements. When paired with our tricistronic Fucci-apoptosis reporters, FluoroFate enables simultaneous monitoring and automated analysis of cell-cycle progression and caspase activation in individual cells. Together, the reporter and analysis platform provide a powerful integrated tool for determining how cell-cycle state influences apoptotic responses to treatments that act at defined stages of the cell cycle. FluoroFate can also classify cells as alive or dead using persistent fluorescent signals and distinguish apoptotic from non-apoptotic cell death based on the temporal ordering of those signals.

## METHODS

### Chemicals and reagents

MDA-MB-231 cells were provided by the Gilmore laboratory (University of Manchester, UK), SKOV3 cells were provided by the Tergaonkar laboratory (A*STAR Institute, Singapore), and Flp-In NIH/3T3 cells were purchased from Thermo Fisher Scientific. SKOV3 RIPK1 KO cells were a gift from Chen Ying in the Tergaonkar lab. Cell lines were maintained in Dulbecco’s Modified Eagle’s Medium (DMEM; Gibco, Thermo Fisher Scientific) supplemented with 10% (v/v) foetal bovine serum (FBS) and no antibiotics. NIH/3T3 cells were maintained in zeocin before targeting and hygromycin after targeting, and this cell line was maintained in Pen/Strep (1%). Dulbecco’s phosphate-buffered saline (DPBS) was purchased from Gibco. Recombinant human tumour necrosis factor alpha (TNFα) was obtained from RayBiotech, birinapant from Apexbio, cycloheximide from Cell Signaling Technology, Annexin V-FITC from Biotium, and propidium iodide from Abcam. All other reagents were purchased from Sigma-Aldrich unless otherwise stated. All cell lines were routinely tested and confirmed to be free of mycoplasma contamination.

### Drug preparation and treatment

Cells were treated with drugs diluted in complete growth medium, at concentrations of 10 ng/ml (TNFα), 0.1 μM (birinapant), 15 μg/ml (cycloheximide), and 1 μM (staurosporine), unless otherwise stated.

### Generation of Fucci-apoptosis reporter constructs

The CA-Cerulean construct was designed based on a previously described caspase-activatable GFP reporter (Nicholls et al., 2011), in which fluorophore maturation is inhibited by a hydrophobic quenching peptide derived from the influenza M2 transmembrane domain. A DEVD caspase recognition sequence was inserted between Cerulean and the quenching peptide, such that caspase 3/7-mediated cleavage releases the fluorophore, allowing proper folding and fluorescence. The construct included a Kozak consensus sequence (GCCACC) upstream of the coding region and was fused at the C-terminus to a P2A self-cleaving peptide (Kim et al., 2011). The CC3AI construct was based on a split-fluorophore design (Zhang et al., 2013), in which the N- and C-terminal fragments of Cerulean were linked via a DEVD caspase cleavage sequence. Upon caspase activation, cleavage enables reconstitution of the fluorescent protein through bimolecular fluorescence complementation. To enhance reconstitution efficiency, split Npu DnaE intein sequences were incorporated to mediate protein trans-splicing (Iwai et al., 2006, Zettler et al., 2009). A Kozak sequence was included at the 5′ end, and the construct was fused to a P2A peptide at the 3′ end.

Both CA-Cerulean and CC3AI sequences were synthesised (GeneArt, Thermo Fisher Scientific) and subcloned upstream of the bicistronic tFucci2(SA) cassette in the pcDNA5-tFucci2(SA) vector (Mort et al 2014). Constructs were inserted via MluI and BssHII restriction sites positioned between the CAG promoter and the mCherry-hCDT1(30/120) component of the tFucci2(SA) system, generating CA-tFucci2(SA) and CC3AI-tFucci2(SA) tricistronic expression vectors. Stable cell lines were generated in NIH 3T3 cells using the Flp-In recombination system (Invitrogen) according to the manufacturer’s protocol, allowing site-specific integration of the constructs to generate an isogenic cell population.

### Cell death assays

#### Flow cytometry

Cells were seeded in 6-well plates (Corning, UK) at a density of 5 × 10⁵ cells per well in 2 ml complete medium and allowed to adhere for 24 h. Cells were treated with TNFα and birinapant. Both floating and adherent cells were collected, pooled, and centrifuged at 400 × g for 10 min at 4°C. Cells were washed with ice-cold PBS and resuspended in 100 μl Annexin V binding buffer. Annexin V-FITC (5 μl) was added, and cells were incubated for 15 min at room temperature in the dark. Following incubation, 400 μl binding buffer containing PI (final concentration 0.25 μg/ml) was added. Samples were analysed using a BD LSRFortessa™ flow cytometer, and data were processed using FlowJo software (BD Biosciences). Data were gated against untreated controls, and 10,000 events were recorded per sample. Live cells were defined as Annexin V-negative/PI-negative.

#### Live-cell imaging (IncuCyte)

Cells were seeded in 12-well plates at a density of 2 × 10⁵ cells per well in 1 ml complete medium and incubated for 24 h prior to treatment. Cells were treated with TNFα, birinapant, Annexin V-FITC (final concentration 1 μg/ml), and PI (0.25 μg/ml final concentration). Images were acquired using an IncuCyte® ZOOM Live-Cell Analysis System at 20× magnification every hour for 24 h. Nine fields per well were imaged. Cells were maintained at 37°C in a humidified atmosphere containing 5% CO₂ throughout imaging.

#### Live-cell imaging (Confocal)

CA-tFucci2(SA) and CC3AI-tFucci2(SA) NIH3T3 cells were seeded into 12-well glass-bottom plates and treated with staurosporine or TNFα (50 ng/ml). Confocal imaging was performed using a Nikon A1R confocal microscope equipped with a resonant scanner and spectral detector (Nikon Eclipse TiE). The following excitation lasers and emission filters were used: Cerulean (457.9 nm excitation, 482/35 emission), mVenus/YFP (514.5 nm excitation, 540/30 emission), and mCherry (561.3 nm excitation, 595/50 emission). Brightfield images were acquired using a transmitted light detector. Live-cell imaging was conducted using an environmental chamber maintained at 37°C with 5% CO₂. Nikon Perfect Focus System was used to minimise focal drift. Images were acquired using NIS-Elements software at 20× magnification every 20 minutes.

### Western blotting

Total protein was extracted using the NucleoSpin RNA/Protein purification kit (Macherey-Nagel) according to the manufacturer’s instructions. Protein samples were mixed with 1× Laemmli buffer (2% SDS, 5% β-mercaptoethanol). Protein concentration was determined using a modified Bradford assay incorporating α-cyclodextrin (5 mg/ml) to reduce SDS interference (Rabilloud, 2018). Proteins were separated on NuPAGE™ 4–12% Bis-Tris gels (Invitrogen) and transferred to PVDF membranes (Bio-Rad). Membranes were blocked with 5% (w/v) skimmed milk in PBST (PBS with 0.1% Tween-20) for 30 min at room temperature and incubated overnight at 4°C with primary antibodies. Membranes were washed and incubated with HRP-conjugated secondary antibodies for 2 h at room temperature. Signals were detected using a Bio-Rad ChemiDoc™ imaging system and quantified using Image Lab software.

The following primary antibodies were used: anti-β-actin (ACTB; Sigma-Aldrich, A2066), anti-cleaved caspase-3 (Epitomics, #1476-S), anti-cleaved caspase-8 (Cell Signaling Technology, #9748S), anti-cIAP1 (Cell Signaling Technology, #7065), anti-PARP (Cell Signaling Technology, #9532S), and anti-phospho-RIPK1 (Ser166) (Cell Signaling Technology, #44590S). The following horseradish peroxidase (HRP)-conjugated secondary antibodies were used: anti-rabbit IgG-HRP (Santa Cruz Biotechnology, sc-2357), and anti-mouse IgG-HRP (Santa Cruz Biotechnology, sc-2005).

### Cell death analysis pipeline

Time-lapse images were analysed using FluoroFate. A breakdown of the FluoroFate analysis pipeline can be found at: https://github.com/isobelth/FluoroFate/blob/main/README.md. Annexin V and PI channels were Gaussian-blurred and thresholded independently to generate binary positive masks. For each tracked cell, overlap between the cell mask and the thresholded fluorescence signal was measured at each frame. Cells were classified as apoptotic when Annexin V positivity preceded PI positivity, and as non-apoptotic when PI positivity preceded or coincided with Annexin V. Cumulative percentages of each fate were then calculated over time at single-cell resolution.

### Statistical tests

Statistical analyses were performed using GraphPad Prism (version 11.0.0). Comparisons between multiple groups were conducted using either one-way ANOVA followed by Dunnett’s post-hoc test (when comparing experimental groups to a single control) or two-way ANOVA (to assess the independent and interactive effects of two variables, such as genotype and treatment). Two-way ANOVA was followed by Fisher’s LSD post-hoc test or Tukey’s multiple comparisons test, depending on whether all pairwise comparisons or a more conservative correction for multiple testing was required.

## ACKNOWLEDGMENTS

We thank Mike White for his exceptional mentorship and support during Marcus K Preedy’s scientific training, John Silke for sharing his expertise in TNFα/TNFR1 signalling, and Huynh Vinh Thang for valuable guidance and discussions.

## AUTHOR CONTRIBUTIONS

Marcus K Preedy: conceptualisation, investigation, methodology, formal analysis, writing of manuscript. Isobel Taylor-Hearn: conceptualisation, methodology, writing review and editing, development of software. Chen Ying: investigation, methodology, supervision. Matthew Ford: conceptualisation, investigation, methodology. Ian J Jackson: funding acquisition, conceptualisation, supervision. Andrew Gilmore: conceptualisation, supervision. Vinay Tergoankar: funding acquisition, supervision. Richard L Mort: funding acquisition, conceptualisation, supervision, writing review and editing.

## DATA AND CODE AVAILABILITY

All data is presented within the paper; all code is available at https://github.com/isobelth/FluoroFate.

## DECLARATION OF GENERATIVE AI AND AI-ASSISTED TECHNOLOGIES

During preparation of this manuscript, the authors used OpenAI Codex (OpenAI) to assist with language editing and development and debugging of ImageJ macros used for image-montage preparation. All AI-assisted text and code were critically reviewed, verified and, where necessary, modified by the authors, who take full responsibility for the final manuscript and analyses. Generative AI was not used to generate primary experimental data.

